# Spectral emission profile and wavelength tolerances affect pulse oximeter performance

**DOI:** 10.64898/2026.04.20.719559

**Authors:** M. Reiser, A. Breidenassel, O. Amft

**Affiliations:** University of Freiburg, Faculty of Engineering, 79110 Freiburg, Germany; HAW Landshut, Faculty of Electrical Engineering, 84036 Landshut, Germany; Hahn-Schickard, 79110 Freiburg, Germany

**Keywords:** Monte Carlo simulation, oxygen saturation, photoplethysmography, pulse oximeter, spectral emission profile, wavelength, wearable

## Abstract

We investigate the effects of skin pigmentation and light source characteristics on the performance of reflective Pulse oximetry (PO) devices used in healthcare and well-being applications. We use Monte Carlo (MC) simulations to compare ideal monochromatic and realistic LED spectral emission profiles and tolerance-related wavelength shifts. The simulation covers photon transport in skin models with melanin concentrations (2.55% to 30.5%) and arterial oxygen saturations SaO_2_ (70% to 100%.) Accuracy was assessed by SpO_2_ error, root-mean-square error RMSE (*A*rms), and percentile tail-errors (P90, P95, and P99).

Monochromatic spectral emission yielded the lowest SpO_2_ error (RMSE = 1.32), while LED spectral emission profiles increased errors (RMSE = 2.10). Infrared wavelength tolerances increased SpO_2_ RMSE by 1.1 ± 0.3. SpO_2_ error increased with melanin concentration, from underestimation (−1.8 ± 0.1%) at 2.55% melanin concentration to overestimation (+3.9 ± 1.2%) at 30.5% for low SaO_2_ (70%) and LED spectral emission profiles. At 30.5% melanin concentration, P95 and P99 exceeded FDA and DIN EN ISO 80601-2-61 thresholds, in particular at low SaO_2_ (70%). Clipping SpO_2_ estimates at 100% resulted in an apparent RMSE decrease of up to 3%, reflecting error masking rather than real error reduction.

In conclusion, LED spectral emission profiles and wavelength tolerances can amplify melanin-related bias in SpO_2_ estimates. Monochromatic emission and tighter wavelength control can reduce SpO_2_ error and should be considered in device design and regulation. Regulatory standards should discourage clipping SpO_2_ estimates at 100% and mandate additional metrics as RMSE fails to reflect clinically critical percentile error thresholds, i.e. P95 and P99.

## Introduction

Photoplethysmography (PPG) is a widely used non-invasive optical method to measure pulsatile changes in blood volume caused by the heartbeat. Reflective PPG measurements are increasingly relevant due to their integration into wearable devices and their ability to continuously monitor physiological parameters (1–3). Pulse Oximetry (PO) is a clinically important application of PPG, which estimates arterial oxygen saturation SaO_2_ by comparing absorption at two wavelengths in the red and infrared spectrum. For PO, light is emitted into tissue and reflections are detected by a photodetector.

The PPG signal consists of two components, the pulsatile (AC) component and the static (DC) component. Static absorbers include skin, bone, and the constant portion of blood volume, while the pulsatile component reflects changes in arterial blood flow. The perfusion index (PI), defined as the ratio of AC and DC components, provides a measure of peripheral perfusion and vascular tone. The ratio of ratios (RoR), defined as the ratio of the PI at the red wavelength to the PI at the infrared wavelength, is calibrated against reference measurements of arterial oxygen saturation SaO_2_.

Skin pigmentation is a critical factor for SpO_2_ performance, as PO tend to overestimate oxygen saturation in individuals with darker skin tones, i.e. increased melanin concentration (4, 5). Skin with increased melanin concentration (i.e., 30.5%) shows a larger absorption coefficient *µ*_*a*_, compared to lighter skin (i.e., 2.55%), with non-linear effects across red and infrared wavelengths. The non-linear signal distortion due to melanin concentration results in a systematic overestimation of oxygen saturation (5), in particular at low levels of arterial oxygen saturation SaO_2_. SpO_2_ estimation error can have serious clinical consequences, in particular when arterial oxygen saturation SaO_2_ is below critical values (e.g., *<* 88% (6, 7)), but the PO device erroneously indicates clinically acceptable levels of SpO_2_ (called occult hypoxemia) (8). The discrepancy in SpO_2_ estimation may delay oxygen therapy or other urgent interventions, especially in darker-skinned patients, who are disproportionately affected. For instance, studies have shown that patients with increased melanin concentration (e.g., 30.5%) are up to three times more likely to experience undetected hypoxemia than patients with low melanin concentration (2.55%) (9).

The accuracy of PO devices is assessed using the root-mean-square error (RMSE, *A*_rms_) between a reference oxygen saturation value SaO_2_ and the SpO_2_ estimation by the PO device. Reference SaO_2_ is obtained through invasive blood gas analysis (10, 11), which is considered as gold standard. However, certified PO devices may still exhibit systematic errors (9, 12, 13), which remain undetected when relying solely on RMSE. Since RMSE only reflects the global average deviation, it does not necessarily capture systematic biases in specific subpopulations, e.g., individuals with increased melanin concentration or low arterial oxygen saturation. In addition, manufacturers often clip SpO_2_ estimations at 100% oxygen saturation, which alters and reduces the subsequently calculated RMSE by masking potential overestimation errors.

For PPG and PO measurements, typically light emitting diodes (LEDs) are used as a light source. LEDs are specified by a nominal wavelength (e.g., 660 nm and 940 nm for PO), usually denoting the centroid of their wavelength emission profile. In reality, LEDs are not monochromatic but emit over a spectral band, e.g., with typical full width at half maximum (FWHM) between 17 nm and 42 nm (SFH 7014C by ams-OSRAM AG). The wavelength spectral emission profile arises from semiconductor band-to-band recombination and thermal carrier distributions. In addition, LEDs are subject to wavelength tolerances, which originate from bandgap variations (including composition, doping, and crystal structure) and manufacturing processes (including epitaxy, layer thickness, defects, and wafer-to-wafer variation). In addition to the intrinsic spectral emission profile, manufacturing variations introduce nominal tolerances ranging from ± 2.5 nm to ± 11.5 nm (e.g., ams-OSRAM SFH 7014C) relative to the specified nominal wavelength. The selection of wavelength is critical in PO because the absorption of oxyhemoglobin O_2_Hb and deoxyhemoglobin HHb change steeply in the red wavelength spectrum (14). Even small shifts can therefore alter the RoR and error in SpO_2_ estimation. Milner and Mathews (15) reported deviations up to 7% for ± 2 nm shifts in the red spectrum, while Rea et al. (16) and Bierman et al. (17) showed that LED spectra contribute to skin-pigmentation bias, which can be mitigated by narrower-band emitters.

Monte Carlo (MC) simulations are a tool to isolate and quantitatively capture complex spectral effects of light sources in PPG and PO (5, 18–20). MC simulations can provide a physically accurate framework to model photon–tissue interactions, including absorption, scattering, and anisotropy, which cannot be captured by simplified Beer–Lambert or diffusion-based approaches. MC simulations can integrate spectral emission profile, detector responsivity, layered and heterogeneous tissue structures (including epidermis, dermis, subcutaneous tissue, bone, or blood vessels), and physiologically relevant changes in blood volume. MC simulations are the gold standard for simulating photon transport in biological media with scattering.

Beyond reproducing the fundamental AC and DC components of the PPG signal, MC simulations offer the chance to analyse photon path length distributions, penetration depth, or PI. Spectral sweeps can be handled efficiently by computing tissue responses across a wavelength grid and applying source and detector spectral weighting in post-processing. Recent studies have successfully employed MC simulations to investigate dual-wavelength PPG formation and the impact of melanin on SpO_2_ estimation error (5, 18, 21).

Milner and Mathews (15) highlighted that nominal wave-length shifts can induce large SpO_2_ estimation errors, yet did not account for spectral emission, scattering, or melanin. Tsiakaka et al. (22) used a simplified absorption model to optimise wavelength pairs, assuming monochromatic sources and homogeneous tissue. Bierman et al. (17) demonstrated experimentally that LED spectral emission increases melanin-dependent SpO_2_ estimation error, but neglected manufacturing tolerances, beam profiles, and additional metrics, including *A*_rms_. In contrast, our study systematically disentangles the independent effects of spectral emission profiles and wavelength tolerances, which to date have either been considered separately or with substantial simplifications.

In this work, we investigate wavelength tolerances and spectral emission profiles of commercially available LEDs to reflect realistic device conditions. Melanin concentrations were selected to represent Fitzpatrick scale skin types (23), and SaO_2_ values were aligned with international PO standards (10, 11). In addition, source-detector distance was varied to disentangle geometry influences from PO performance. Empirical SpO_2_ estimations are sensitive to physiological variability and other non-controllable parameters, including peripheral perfusion of individuals and temperature-dependent wavelength shifts. By employing MC simulations, we incorporate realistic tissue heterogeneity (epidermis, dermis, subcutaneous fat, muscle), melanin concentrations, detector responsivity, and photon scattering, modelling a physical representation of the two physiological phases (i.e. diastole and systole) in PPG formation. Unlike previous work, we quantify the spectral effects not only via absolute SpO_2_ estimation error (RMSE, *A*_rms_), but analyse the RMS deviation of SpO_2_ estimation as well as tail-errors. Furthermore, by analysing wavelength tolerance and spectral emission profile under identical simulation conditions, we provide the first direct comparison of their relative impact. Our approach allows us to derive actionable design insights for light source specifications, source–detector geometry, and PO device calibration strategies that remained unaddressed so far.

In summary, this paper provides the following contributions:

1. We perform realistic MC simulations using a validated framework to analyse SpO_2_ estimation performance in wearable and mobile health devices. Our analysis covers tissue heterogeneity, melanin concentrations, physiological phases, detector responsivity (spectral and angular), light source spectral and angular emission profiles, nominal wavelength tolerance, and source-detector distance variability, to reflect conditions encountered in wearable SpO_2_ monitoring.
2. We introduce an extended error analysis beyond absolute SpO_2_ estimation error that includes tail-error quantification.
3. We derive actionable design insights for light source specifications, source-detector geometries, and calibration strategies for pulse oximeters.

## Methods and Materials

We deployed a framework consisting of three main components: (1) combined skin and sensor model, (2) photon-skin simulation, and (3) reflective PO simulation (see Fig. 1). The skin model included layer-specific optical characteristics (absorption coefficient *µ*_*a*_, scattering coefficient *µ*_*s*_, anisotropy factor *g*, and refractive index *n*), anatomical characteristics (skin layers, thickness), melanin concentration, arterial oxygen saturation SaO_2_, and physiological states (systole and diastole). The sensor model included the spectral and angular sensitivity of the photodiode and angular and spectral emission profile of both the LED and an assumed ideal monochromatic light source. The photon-skin simulation propagated photon packets, based on the sensor model, through our skin model. For each configuration, we recorded the detected intensity *I*, photon packet count, absorption per skin layer, and computation time across source-detector distances ranging from 3.5 mm to 11.5 mm.

**Fig. 1.**
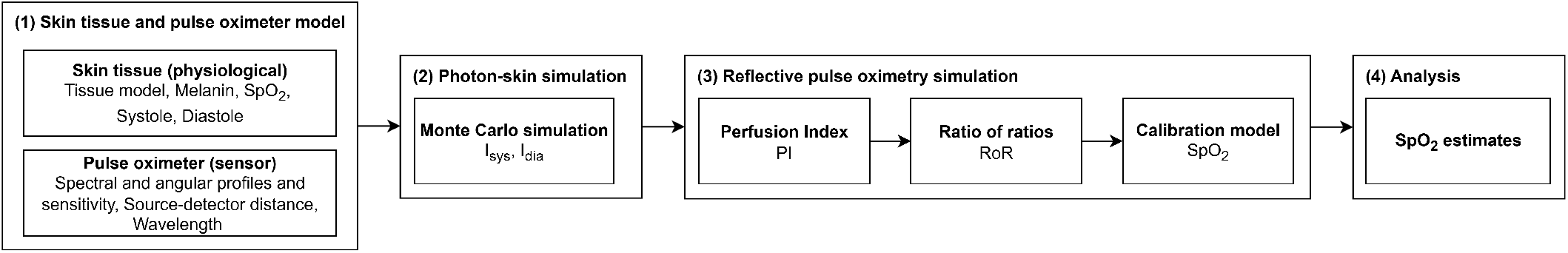
Overview of our simulation approach, comprising (1) skin tissue and pulse oximeter modelling (skin physiology and sensor configuration), (2) photon–skin simulation using Monte Carlo (MC) methods, (3) reflective pulse oximetry (PO) simulation to generate perfusion index (PI), ratio of ratios (RoR), SpO2 calibration constants (A,B,C), and (4) analysis using root-mean-square error of SpO_2_ estimation (RMSE, i.e. average RMSE *A*_rms_) and RMSE distribution tail-errors (P90, P95, P99).

### A. Sensor model

As a light source, we modelled the LED SFH7014C and as a detector the photodiode SFH2704A (both by ams-OSRAM AG). The spatial angular emission profile was represented by the relative source intensity *I*_rel_ (see Fig. 2).

**Fig. 2.**
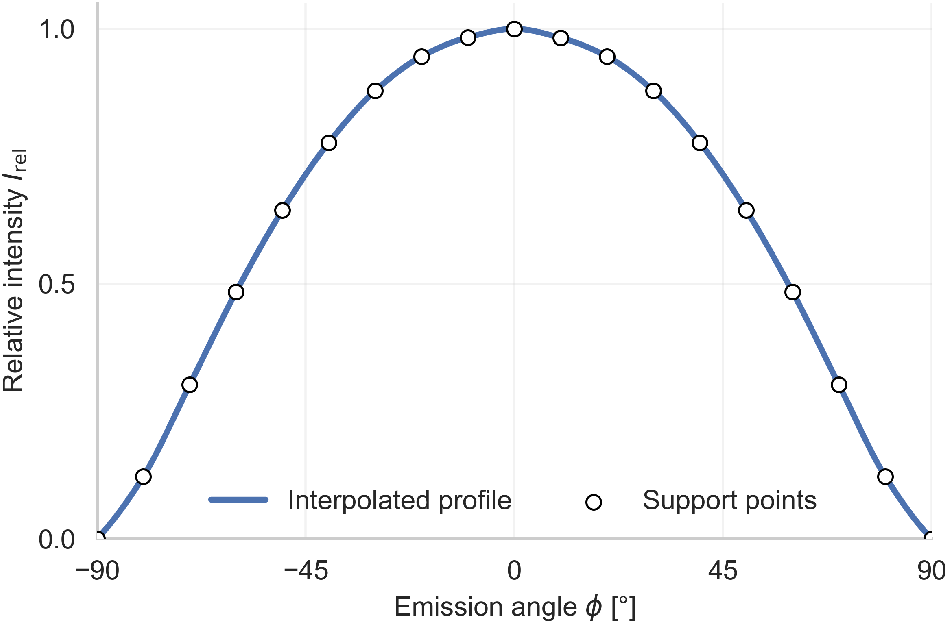
Spatial angular emission profile of an exemplary LED (SFH7014C by OSRAM) shown as relative radiant intensity *I*_rel_ across emission angleφ. Support points ranged from −90°to +90°in steps of 10°.

For tolerance analysis, we varied the nominal wavelengths within their specified tolerance ranges, i.e., 655 nm ± 2.5 nm for the red spectrum and 940 nm ± 9.5 nm for the infrared spectrum. For LED spectral emission simulations, we simulated spectral emission at FWHM in the red spectrum of 17 nm and in the infrared spectrum of 42 nm. Tab. 1 shows the relative spectral emission *I*_e,rel_ defined at discrete support points. For spectral emission simulations of the monochromatic light source, we assumed an ideal spectral emission profile and simulated the nominal wavelength in the red and infrared spectrum (655 nm and 940 nm).

**Table 1.** Relative spectral emission *I*_e,rel_ with support points for the red (655 nm) and infrared (940 nm) LEDs.

| $I_{\text{e,rel}}$ | Wavelength [nm] | |
| --- | --- | --- |
| | $\lambda_{\text{R}}$ (655) | $\lambda_{\text{IR}}$ (940) |
| 0.2 | 642 | 900 |
| 0.4 | 649 | 919 |
| 0.6 | 653 | 930 |
| 0.8 | 657 | 938 |
| 1.0 | 662 | 950 |
| 0.8 | 665 | 959 |
| 0.6 | 667 | 963 |
| 0.4 | 670 | 968 |
| 0.2 | 673 | 975 |

The detected Intensity *I* was adjusted to the relative spectral sensitivity of the photodiode *S*_S,*λ*_ and the directional characteristics *S*_rel,*φ*_.

### B. Skin model and photon–skin simulation

We used a previously developed multilayer skin model which consists of six skin layers (*epidermis, capillary loops, upper plexus, reticular dermis, deep plexus*, and *hypodermis*), and an additional *muscle* layer (5). We simulated both physiological states, systole and diastole. During systole, the blood volume fraction was increased in skin layers (*capillary loops, upper plexus, reticular dermis, deep plexus*, and *hypodermis*) (19). The photon-skin simulations were performed using a previously developed MC framework for photon-tissue interactions, implemented in C++ and CUDA (24). For each parameter configuration (defined by physiological state, wave-length *λ*, monochromatic or LED spectral emission profile, melanin concentration *C*_Mel_, and arterial oxygen saturation SaO_2_) a total of 5 *×* 109 photon packets were simulated. Each configuration required an average of 234.1 ± 22.8 s of processing on an NVIDIA RTX 6000 Ada.

### C. Reflective pulse oximetry simulation

Oxygen is transported bound to hemoglobin in erythrocytes. The arterial oxygen saturation SaO_2_ is defined as the fraction of O_2_Hb relative to the total functional hemoglobin (O_2_Hb + HHb). Hemoglobin absorbs light differently depending on its oxygenation state. The absorption coefficient *µ*_*a*_ of O_2_Hb is greater in the infrared wavelength spectrum compared to the red wavelength spectrum, with approximately 4.3-fold higher absorption at 940 nm compared to 655 nm. (14). In contrast, HHb increases absorption in the red wavelength spectrum compared to the infrared spectrum, with approximately 3.8-fold greater absorption at 655 nm compared to 940 nm (14). The opposing absorption properties of O_2_Hb and HHb in the red and infrared spectrum can be used to empirically estimate oxygen saturation SpO_2_.

For estimating SpO_2_, PI was derived as the ratio of the pulsatile to the non-pulsatile signal components:

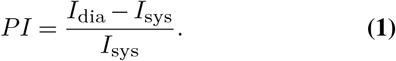

Based on the PI in the red and infrared spectrum, RoR was calculated as the relative relationship between the distinct absorption characteristics of O_2_Hb and HHb:

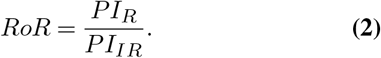

Using a manufacturer-specific empirical calibration, the arterial oxygen saturation SaO_2_ was estimated from RoR as:

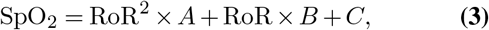

where A, B, and C are calibration constants.

PO devices can be classified as medical devices and thus their use is regulated by international standards, including DIN EN ISO 80601-2-61(11), and the U.S. Food and Drug Administration (FDA) (10). To ensure reliability, the regulations specify minimum performance requirements. RMSE, denoted as *A*_rms_, quantifies the deviation between the reference oxygen saturation *S*_R_ (e.g., determined by blood gas analysis) and the oxygen saturation SpO_2_ estimated by the PO. The RMSE is calculated as:

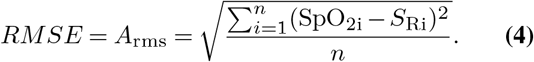

The RMSE limit required by the FDA for reflective PO is below 3.5% and by DIN EN ISO 80601-2-61 below 4.0%. For each parameter configuration (SaO_2_, systolic and diastolic state, melanin concentration, source-detector distance, wavelength combination, and monochromatic or LED spectral emission profile), we performed simulations with 25 seeds with 5 *×* 109 photon packets each. The parameter seeds were used to introduce variability in photon interactions. Different seeds resulted in variations in simulation outcomes (including photon path length and scattering angle). We interpreted the parameter seed as natural interpersonal variability, corresponding to a virtual study cohort.

Based on the virtual study cohort, a general calibration curve was calculated. The deviation between the estimated oxygen saturation SpO_2_ and the actual input simulation parameter for oxygen saturation (denoted as SaO_2_ or *S*_R_) was then evaluated to determine the RMSE. The calibration and RMSE calculation was repeated for all simulated source–detector distances ranging from 3.5 mm to 11.5 mm.

### D. Experiments and validation

In this study, we investigated the impact of two hardware-related factors on reflective PO: (1) manufacturing-related tolerances in nominal LED wavelengths and (2) the spectral emission profile of an LED compared to an assumed ideal monochromatic light source. To quantify the impact in both cases, MC simulations were performed for each scenario. For tolerance analysis, we simulated the endpoints of the specified wavelength ranges (*±* 2.5 nm at 655 nm and ± 9.5 nm at 940 nm). For spectral emission, both LED and ideal monochromatic light sources were modelled. The approach allowed us to disentangle the independent and combined contributions of tolerances and spectral emission profiles under otherwise identical conditions. The MC framework was validated against laboratory measurements using a porcine skin phantom (24, 25) with angular resolution on both beam incidence and detector detection angles. Furthermore, we validated the MC framework against real-world PPG measurements of systolic and diastolic signal levels from a participant study (26).

The resulting RoR values were calibrated against the reference oxygen saturation (arterial oxygen saturation SaO_2_ of the skin model), and the estimation accuracy was quantified using the RMSE. The statistical significance of SpO_2_ estimation analyses was assessed using the paired Wilcoxon signed-rank test (*α* = 0.05 and *α* = 0.01), comparing (1) tolerance-related wavelength shifts relative to the 655/940 nm reference and (2) monochromatic versus LED spectral emission profiles. In addition to regulation imposed by the international standards, we analysed tail error quantiles (P90, P95, and P99) of absolute SpO_2_ estimation error to capture clinically critical deviations in the tails of the error distribution. Increased melanin concentration can systematically influence reflected signal intensities in the red and infrared spectrum, leading to a broader and potentially skewed error distribution in SpO_2_ estimation. In consequence, individuals with higher melanin levels are more likely to be represented in the extreme error tails rather than by a single deterministic error value.

The analysis was restricted to source-detector distances, representing near-optimal sensor configurations (3.5 mm and 4.5 mm) as a wider range of distances would introduce geometric effects, i.e. alter optimal photon path, which obscure the effects of the investigated variables (i.e., spectral emission profiles, wavelength tolerance).

## Results

### E. Wavelength tolerance

Fig. 3 shows the SpO_2_ estimation RMSE for all wavelength tolerance combinations of the red spectrum (652.5 nm, 655 nm, 675.5 nm) and the infrared spectrum (930.5 nm, 940 nm, 949.5 nm). PO calibration was performed based on nominal wavelengths 655 nm and 940 nm, respectively. RMSE increased with source-detector distance. In the infrared spectrum, wavelengths shifts towards 949.5 nm showed worst RMSE compared to shifts towards 930.5 nm. Lowest RMSE was achieved for source-detector distances of 3.5 mm and 4.5 mm.

**Fig. 3.**
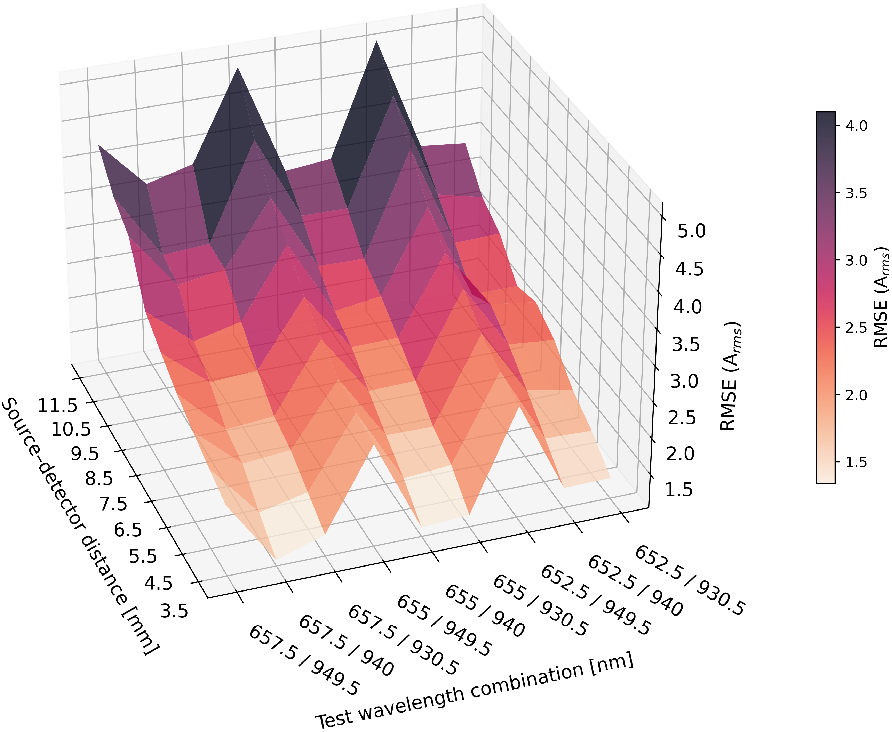
RMSE (*A*_rms_) of wavelength tolerance combinations for SpO_2_ estimation considering source-detector distance. Wavelength in the red spectrum was 655 nm ± 2.5 nm and in the infrared spectrum 940 nm ± 9.5 nm (calibration on 655 nm and 940 nm, respectively). RMSE increased with increasing source-detector distance and varies with wavelength combinations in the red and infrared spectrum.

Fig. 4 shows the SpO_2_ estimation error (i.e., SaO_2_ − SpO_2_) depending on melanin concentration. Here, all wavelength tolerance combinations for both calibrations (655 nm and 940 nm) as well as all SpO_2_ levels, seeds, and source-detector distances were are aggregated. Lower skin pigmentation (2.55% melanin concentration), yielded an average error of −2.15 ± 1.64%, moderate skin pigmentation (15.5% melanin concentration) an average of −0.56 ± 1.73%, and higher skin pigmentation (30.5% melanin concentration) an average of 1.21 ± 3.31%. With increasing melanin concentration, the average SpO_2_ estimation error shifts from underestimation to overestimation. Distribution shift in Fig. 4A was caused by the varying source–detector distances. The distribution shift is most pronounced at 2.55% melanin due to higher SNR at low melanin concentrations (27), which helps to resolve distance-dependent differences. With increasing melanin concentrations, increased absorption reduces SNR that blurs the effect of varying source–detector distances.

**Fig. 4.**
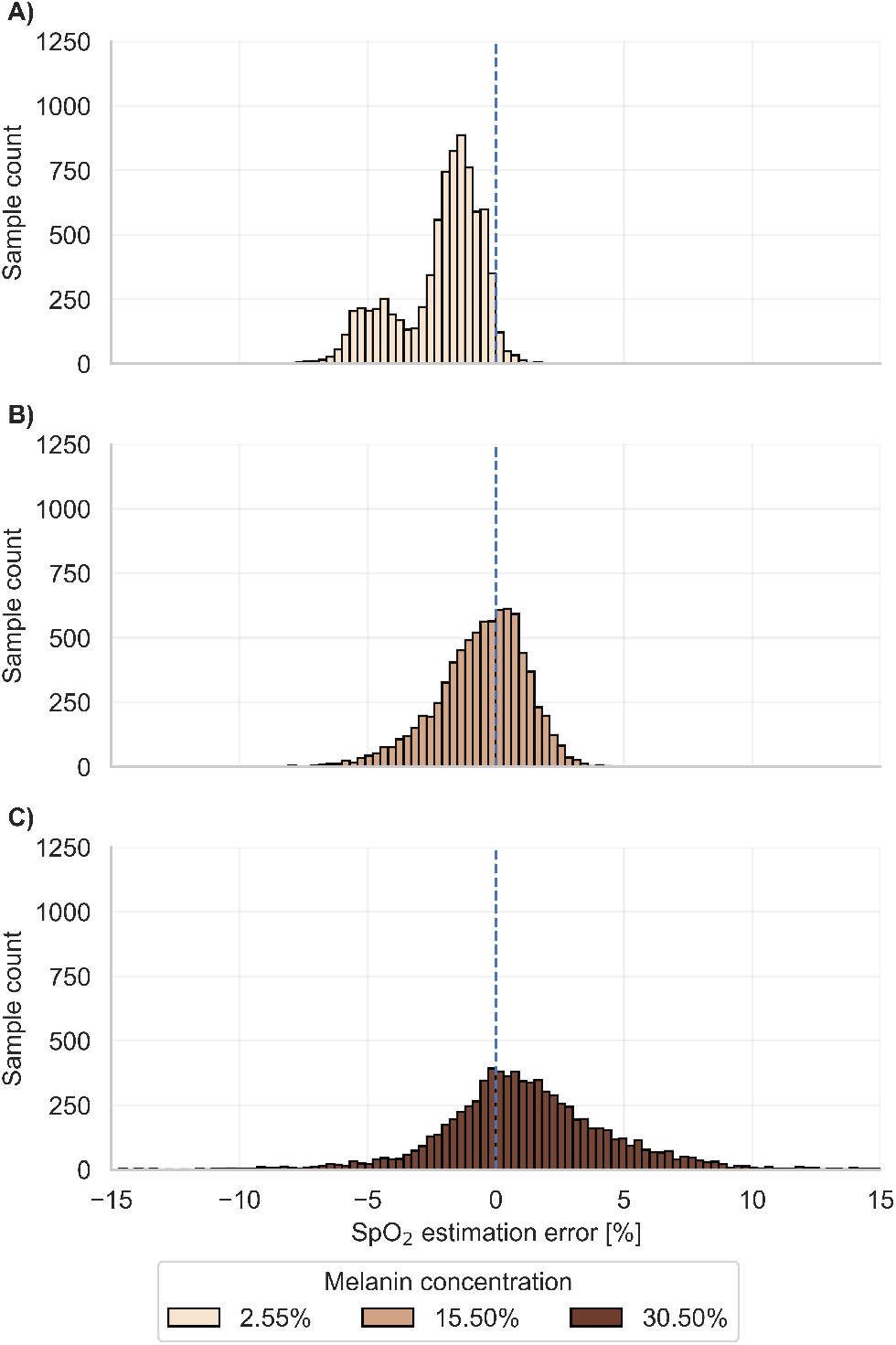
Distribution of SpO_2_ estimation error (SaO_2_ − SpO_2_) for individual melanin concentrations (A: 2.55%, B: 15.5%, C: 30.5%). Here, all wavelength tolerance combinations for both calibrations (655 nm and 940 nm) as well as all SpO_2_ levels, seeds, and source-detector distances of 3.5 mm and 4.5 mm are aggregated.

Wavelength combinations of 657.5/940 nm yielded the lowest RMSE of 1.26, whereas 652.5/949.5 nm resulted in the highest RMSE of 2.64 (see Fig. 5). When infrared wave-length shifted to 949.5 nm, the highest average increase in RMSE of 1.04 ± 0.20 was observed. All deviations relative to 655 nm/940 nm were statistically significant, except for 657.5 nm/940 nm (see Tab. 2).

**Fig. 5.**
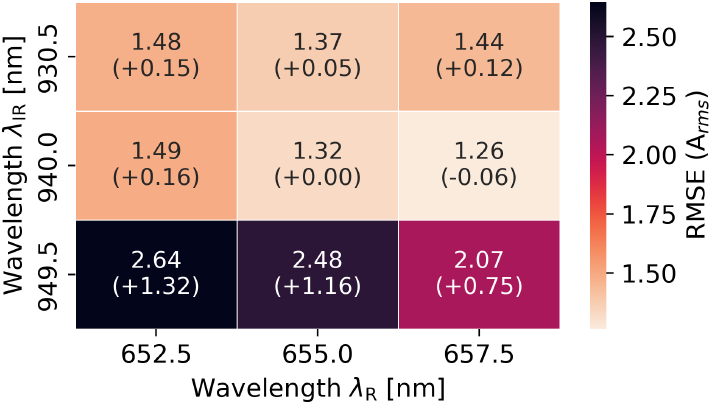
RMSE (*A*_rms_) of SpO_2_ estimation for wavelength tolerance combinations (655 nm ± 2.5 nm and 940 nm ± 9.5 nm) with corresponding calibration at 655 nm and 940 nm. Results were aggregated over all SpO_2_ levels, seeds, and source-detector distances of 3.5 mm and ± 4.5 mm. Numbers: *A*_rms_, [Δ*A*_rms_], where Δ*A*_rms_ = *A*_rms(*λ*R,*λ*IR)_ − *A*_rms(655 nm,940 nm)_.

**Table 2.** Wilcoxon signed-rank results to test with Holm correction for statistical significant differences between RMSE (A_rms_) comparisons across manufacturer-related wavelength tolerances.

| Hypothesis | corrected $p$-value |
| --- | --- |
| 652/930 vs 655/940 | $1.33 \times 10^{-2} *$ |
| 652/940 vs 655/940 | $< 1.0 \times 10^{-3} **$ |
| 652/950 vs 655/940 | $< 1.0 \times 10^{-3} **$ |
| 655/930 vs 655/940 | $1.57 \times 10^{-2} *$ |
| 655/950 vs 655/940 | $< 1.0 \times 10^{-3} **$ |
| 657/930 vs 655/940 | $3.06 \times 10^{-2} *$ |
| 657/940 vs 655/940 | $1.28 \times 10^{-1}$ |
| 657/950 vs 655/940 | $< 1.0 \times 10^{-3} **$ |

Corrected $p$ -values were obtained using the Holm method. Significance levels: \* $p < 0.05$ , \*\* $p < 0.01$ .

### F. Spectral emission profile

Fig. 6 compares simulated oxygen saturation SpO_2_ estimates and reference arterial oxygen saturation SaO_2_. Both, LED and monochromatic spectral emission profiles showed an increasing SpO_2_ overestimation for high melanin concentrations (30.5%) and oxygen saturation level SaO_2_ ≤ 80% (see Fig. 6A and B). The monochromatic profile provided more accurate SpO_2_ estimates (RMSE: 1.32) compared to simulations with the LED profile (RMSE: 2.10). Clipping SpO_2_ estimations at 100% resulted in systematic overestimation and an apparent reduction of RMSE by 2.9% for the LED profile and by 3.0% for the monochromatic profile (see Fig. 6C and D). All comparisons between the LED and monochromatic emission profiles were statistically significant (see Tab. 3).

**Fig. 6.**
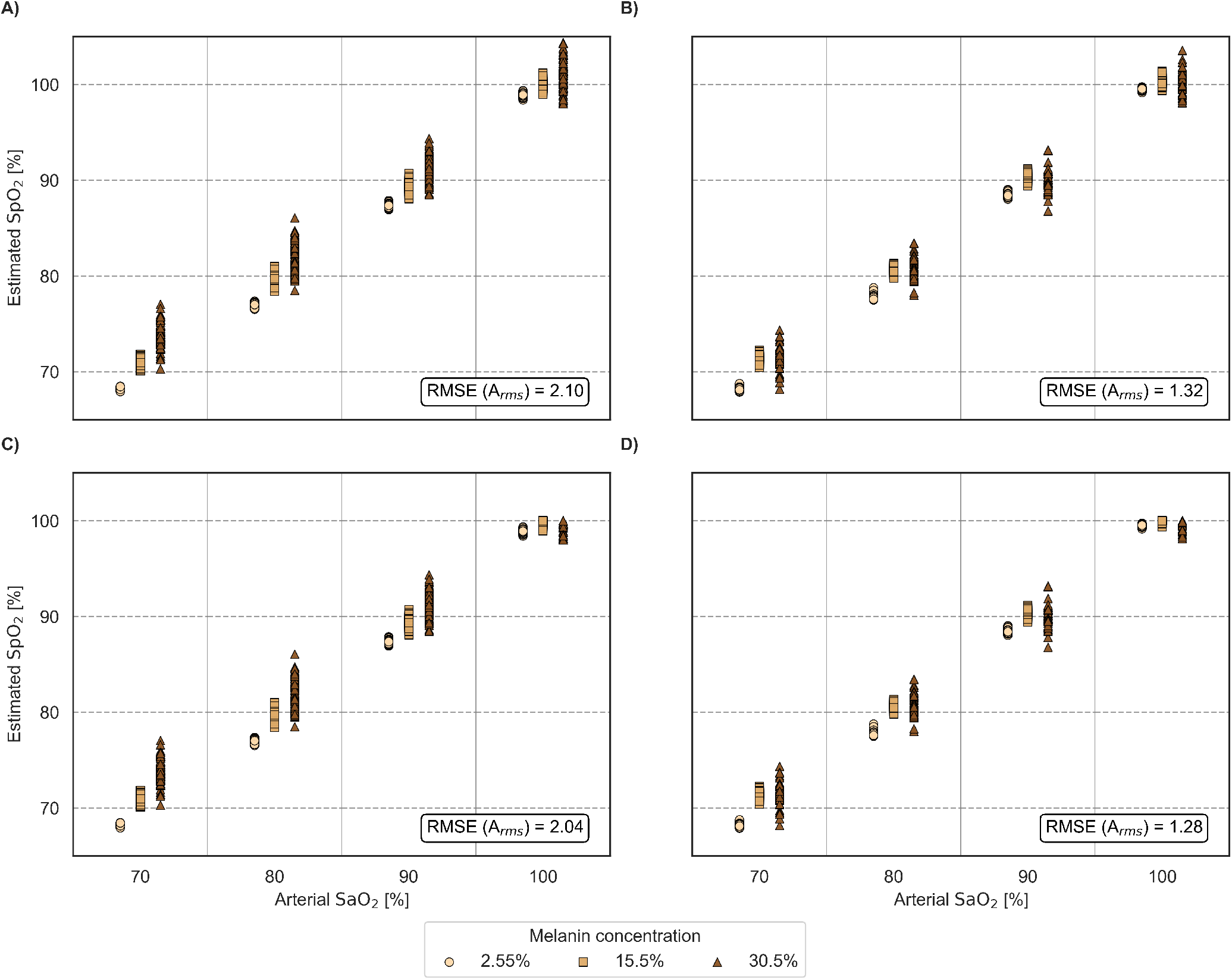
Comparison of simulated oxygen saturation SpO_2_ estimates and reference oxygen saturation SaO_2_ for spectral emission profiles (A, C: LED emission spectrum, B, D: monochromatic emission spectrum) for source-detector distances of 3.5 mm and 4.5 mm. In addition the effect of clipping at 100% oxygen saturation is compared (A, B: no clipping; C, D: clipping). Each symbol denotes a different melanin concentration *C*_*Mel*_. Dashed lines indicate reference SaO_2_ values.

**Table 3.** Overview of deviations (SpO_2_ − SaO_2_) per melanin concentration and spectral emission profiles (LED, monochromatic). Results are reported as median [Q1/Q3]. Statistical significance is assessed using Wilcoxon signed-rank *p*-values with Holm correction for multiple comparisons.

| SaO<sub>2</sub><br>[%] | Melanin<br>[%] | LED | Monochromatic | corrected *p*-value |
| --- | --- | --- | --- | --- |
|  |  | median [Q1/Q3] [%] |  |  |
| 70 | 2.55 | -1.78 [-1.87/-1.69] | -1.66 [-1.80/-1.50] | < 1.0 × 10<sup>-3</sup> ** |
|  | 15.5 | +1.03 [+0.74/+1.24] | +1.41 [+1.19/+1.61] | < 1.0 × 10<sup>-3</sup> ** |
|  | 30.5 | +3.88 [+3.33/+4.66] | +1.58 [+0.97/+1.99] | < 1.0 × 10<sup>-3</sup> ** |
| 80 | 2.55 | -2.95 [-3.08/-2.85] | -1.92 [-2.17/-1.59] | < 1.0 × 10<sup>-3</sup> ** |
|  | 15.5 | -0.37 [-0.68/-0.14] | +0.71 [+0.50/+0.94] | < 1.0 × 10<sup>-3</sup> ** |
|  | 30.5 | +2.01 [+1.30/+3.11] | +0.82 [+0.09/+1.71] | 1.15 × 10<sup>-3</sup> ** |
| 90 | 2.55 | -2.59 [-2.73/-2.51] | -1.41 [-1.69/-1.24] | < 1.0 × 10<sup>-3</sup> ** |
|  | 15.5 | -0.65 [-0.91/-0.32] | +0.34 [+0.05/+0.63] | < 1.0 × 10<sup>-3</sup> ** |
|  | 30.5 | +1.47 [+0.44/+2.39] | +0.13 [-0.42/+0.83] | < 1.0 × 10<sup>-3</sup> ** |
| 100 | 2.55 | -1.08 [-1.22/-0.94] | -0.52 [-0.56/-0.43] | < 1.0 × 10<sup>-3</sup> ** |
|  | 15.5 | -0.04 [-0.28/+0.20] | +0.32 [+0.04/+0.56] | < 1.0 × 10<sup>-3</sup> ** |
|  | 30.5 | +1.05 [+0.02/+1.85] | +0.02 [-0.81/+1.14] | 1.11 × 10<sup>-2</sup> * |

Corrected $p$ -values were obtained using the Holm method. Significance levels: \* $p < 0.05$ , \*\* $p < 0.01$ .

Fig. 7 shows absolute SpO_2_ estimation error distributions quantiles (P90, P95, P99) per melanin concentration and arterial oxygen saturation SaO_2_. For 2.55% and 15.5% melanin concentration, P90 and P95 remained below the FDA and DIN EN ISO 80601-2-61 thresholds at all oxygen saturation levels. At 30.5% melanin concentration, P95 and P99 frequently exceeded the regulatory limits, in particular at oxygen saturations SaO_2_ ≤ 80%.

**Fig. 7.**
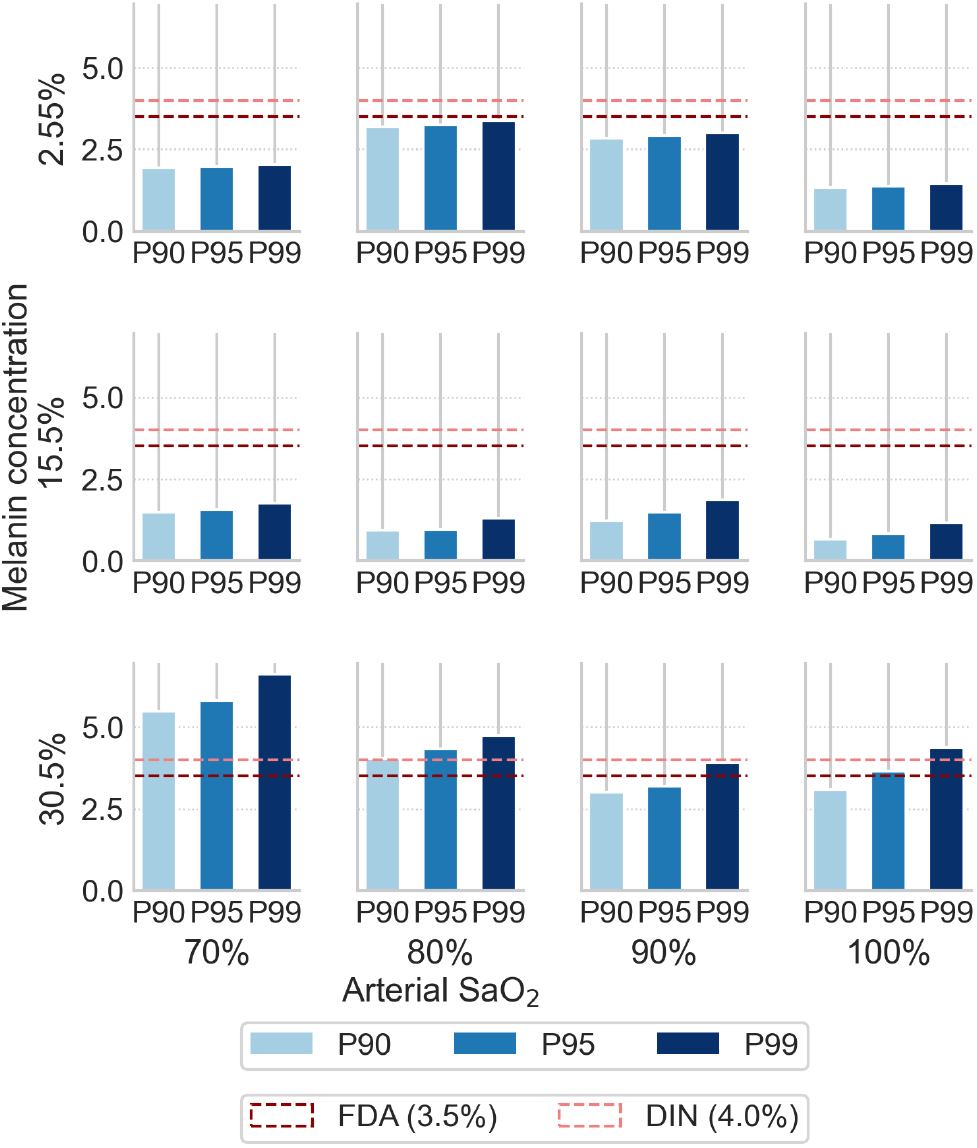
Tail error quantiles (P90, P95, P99) of absolute SpO_2_ estimation error stratified by melanin concentration (rows) and arterial oxygen saturation SaO_2_ (columns). Regulatory limits from FDA (3.5%) and DIN EN ISO 80601-2-61 (4.0%) are illustrated.

## Discussion

Our MC simulation approach allowed us to analyse light source emission profiles and wavelength tolerances under otherwise identical conditions, i.e., without confounding influences (e.g., sensor misalignment, day-to-day physiological variability, and measurement noise). The analysis is vital to guide PO device design considering different skin tones.

Source-detector distances of 3.5 mm and 4.5 mm showed the lowest RMSE. With increasing source-detector distance, photons penetrate deeper tissue layers with larger blood volume fractions and spectral overlap effects (e.g., difference in absorption coefficients *µ*_*a*_ of O_2_Hb and HHb) become more pronounced.

Simulations of wavelength tolerances showed that deviations from the nominal wavelength increased the SpO_2_ estimation error, with RMSE increasing up to 100% relative to the nominal wavelength configuration (see Fig. 5). In our simulations, the wavelength shift from 940 nm to 949.5 nm resulted in the largest RMSE increase from 1.32 to 2.64, primarily due to the increased wavelength-dependent differences in the absorption coefficient of O_2_Hb and HHb (14). In addition, the increased spectral sensitivity of the photodiode in the infrared range further amplifies the infrared contribution compared to the red wavelength in wavelength tolerance shifts.

Wavelength shifts in the red and infrared ranges are critical, as they can alter the resulting RoR to the extent that RMSE exceeds regulatory limits (FDA and DIN EN ISO 80601-2-61) (see Fig. 3). Therefore, light source and photodiode should be selected with minimal wavelength tolerances. For the light source, it is essential to ensure that wavelength-dependent absorption differences between O_2_Hb and HHb remain as small as possible in the relevant wavelength regions. For the photodiode, the tolerance-related difference in wavelength-dependent spectral sensitivity should be minimised. Future studies may explore wavelength combinations in the red and infrared spectral range that are minimally influenced by spectral emission characteristics or wavelength tolerances.

Simulated LED spectral emission (FWHM of 17 nm in the red and 42 nm in the infrared spectrum) resulted in an average increase in RMSE of 59.1% compared to monochromatic spectral emission (see Fig.6). The effects of the LED spectral emission profile amplified the dependence of SpO_2_ estimation performance on melanin concentration, resulting in increased overestimation of SpO_2_ at higher skin pigmentation (30.5% melanin concentration). In particular, SpO_2_ overestimation is clinically relevant and can lead to delayed diagnosis and treatment of individuals.

The influence of tolerance-related wavelength shifts and spectral emission profiles on SpO_2_ estimation are not independent, resulting in a cumulative and asymmetric error distribution (see Fig. 5, 6, and 7). Wavelength tolerances and emission profiles should be restricted to spectral regions, where absorption differences between O_2_Hb and HHb are minimised. In particular, in the red region around 700 nm and in the infrared region around 920 nm, even minor deviations could substantially amplify SpO_2_ estimation error.

Our simulations demonstrate that even small wavelength shifts of 2.5 nm can lead to clinically relevant SpO_2_ estimation bias. Prior work shows that wavelength deviations of 4 nm can lead to an SpO_2_ estimation error of 2% to 7%, increasing with decreasing oxygen saturation (15). Our results are consistent with literature and further show that wavelength-dependent errors persist under realistic conditions, including spectral emission profiles, varying melanin concentrations, and varying source-detector geometries. We systematically analysed and directly compared the spectral emission profile of light sources and wavelength tolerance shifts under identical conditions. For the first time, we assessed wavelength tolerance shifts and LED spectral emission using RMSE, tail-error thresholds (P90, P95, P99), and connected the results to regulatory requirements (FDA and DIN EN ISO 80601-2-61). Our simulations underline that hardware-level design choices directly influence health equity: The same device may perform within acceptable limits in individuals with light skin tone (e.g., 2.55% melanin concentration), but fail disproportionately in individuals with higher melanin concentrations (e.g., 30.5%). Our results confirm that in order to optimise SpO_2_ estimation, melanin range calibration might benefit individuals with increasing melanin concentration (5, 28).

SpO_2_ estimation is influenced by numerous parameters, including beam incidence angle, beam profile, and calibration. In addition, the broad spectral emission profile of LEDs and typical wavelength tolerances can contribute to an overestimation of SpO_2_ estimation in individuals with increased skin pigmentation (e.g., 30.5% melanin concentration), thereby increasing the risk of occult hypoxemia. The systematic error in SpO_2_ estimation can contribute to delays in oxygen therapy or inaccurate clinical assessments. Our findings indicate that hardware specifications can help to reduce inequities: (1) narrow emission profile or monochromatic light sources, and (2) more rigorous selection and binning of light sources with respect to wavelength tolerances reduce SpO_2_ estimation error.

Beyond current standards, which primarily define aggregated error metrics, including RMSE, our findings argue for more explicit spectral specifications in regulations. In particular, maximum allowable FWHM and wavelength tolerances should be required in approval processes, especially for devices intended for a population with varying skin tone. From an ethical perspective, overlooking hardware-level induced errors risk reinforcing disparities in healthcare delivery, in particular in ubiquitous systems, where adoption spans large and heterogeneous user groups.

Furthermore, we highlighted the critical limitation of relying solely on RMSE when evaluating PO performance. While RMSE may suggest compliance to literature, the extreme error tails (see Fig. 7) reveal systematic underperformance with increasing melanin concentration. Exceeding of FDA and DIN EN ISO 80601-2-61 thresholds in P95 and P99 implies that up to 5% of SpO_2_ estimations could be clinically unreliable in individuals with increased skin pigmentation, in particular at low oxygenation. We conclude that (1) devices that meet the global RMSE criteria may still fail disproportionately in populations with increased melanin concentrations, and (2) reporting percentile error thresholds (e.g., P95 and P99) stratified by skin pigmentation may be necessary to ensure safety in diverse populations. In addition, regulatory standards should prohibit the clipping of SpO_2_ estimations at 100% as it masks systematic overestimation and artificially improves the apparent performance of PO devices.

While our MC simulations provide a controlled and reproducible setting, they inevitably abstract real-world conditions. The skin model was static and layered, excluding interindividual variability in skin layer thickness, melanin distribution, or vascular density. Motion artefacts, temperature-dependent wavelength drifts, and sensor placement variability were not included, although they are critical in everyday use of wearable devices. However, precisely the abstraction of our simulations highlight the isolated contribution of spectral emission profile and wavelength tolerances, which are difficult to disentangle in-vivo.

## Conclusions

We demonstrated that both wavelength tolerances and spectral emission profiles significantly affect SpO_2_ estimation accuracy (RMSE). SpO_2_ estimation error increased with increasing melanin concentration, leading to systematic overestimation when arterial oxygen saturation SaO_2_ declined and thus increasing the risk of occult hypoxemia.

SpO_2_ estimation errors due to wavelength tolerance shifts and LED spectral emission profiles were additive and in combination could exceed FDA and DIN EN ISO 80601-2-61 regulatory thresholds. For PO device design, narrow-band LEDs or alternative monochromatic light sources combined with minimal wavelength tolerances could reduce SpO_2_ estimation error and mitigate disparities.

Current regulatory standards rely on RMSE and therefore miss critical tail errors. Percentile error thresholds (P95 and P99) should be considered to properly assess device performance across diverse populations. Furthermore, clipping of SpO_2_ estimates at 100% may mask systematic overestimation and should be discouraged. Detailed specifications of the sensors used in PO devices (i.e., FWHM, wavelength tolerances) are encouraged.

Overall, PO device accuracy is a multi-parameter optimisation problem (including wavelength tolerances, spectral emission profile, PO calibration strategies, and sensor geometry). Addressing all interacting factors is essential for a consistent PO device performance.

## ACKNOWLEDGEMENTS

This work has been submitted to the IEEE for possible publication. Copyright may be transferred without notice, after which this version may no longer be accessible. This work was supported by the Federal Ministry for Economic Affairs and Climate Action (BMWK) on the basis of a decision by the German Bundestag.

## Notes

### Competing Interest Statement

The authors have declared no competing interest.

## Bibliography

1. Mohsen Masoumian Hosseini, Seyedeh Toktam Masoumian Hosseini, Karim Qayumi, Shahriar Hosseinzadeh, and Seyedeh Saba Sajadi Tabar. Smartwatches in healthcare medicine: assistance and monitoring; a scoping review. BMC Medical Informatics and Decision Making, 23(1):248, November 2023. ISSN 1472-6947. doi: 10.1186/s12911-023-02350-w.

2. Manting Chen, Hailiang Wang, Lisha Yu, Eric Hiu Kwong Yeung, Jiajia Luo, Kwok-Leung Tsui, and Yang Zhao. A Systematic Review of Wearable Sensor-Based Technologies for Fall Risk Assessment in Older Adults. Sensors, 22(18):6752, September 2022. ISSN 1424-8220. doi: 10.3390/s22186752.

3. Emma Capulli, Ylenia Druda, Francesco Palmese, Abdul Haleem Butt, Marco Domenicali, Anna Giulia Macchiarelli, Alessandro Silvani, Giorgio Bedogni, and Francesca Ingravallo. Ethical and legal implications of health monitoring wearable devices: A scoping review. Social Science & Medicine, 370:117685, April 2025. ISSN 02779536. doi: 10.1016/j.socscimed.2025.117685.

4. Philip E. Bickler, John R. Feiner, and John W. Severinghaus. Effects of Skin Pigmentation on Pulse Oximeter Accuracy at Low Saturation. Anesthesiology, 102(4):715–719, April 2005. ISSN 0003-3022. doi: 10.1097/00000542-200504000-00004.

5. Maximilian Reiser, Oliver Amft, and Andreas Breidenassel. Analysis of Melanin Concentration on Reflective Pulse Oximetry Using Monte Carlo Simulations. IEEE Access, 13: 24454–24462, 2025. ISSN 2169-3536. doi: 10.1109/ACCESS.2025.3538281.

6. Christopher F Chesley, Meghan B Lane-Fall, Venkat Panchanadam, Michael O Harhay, Arshad A Wani, Mark E Mikkelsen, and Barry D Fuchs. Racial Disparities in Occult Hypoxemia and Clinically Based Mitigation Strategies to Apply in Advance of Technological Advancements. Respiratory Care, 67(12):1499–1507, December 2022. ISSN 0020-1324. doi: 10.4187/respcare.09769.

7. Ashraf Fawzy, Tianshi David Wu, Kunbo Wang, Matthew L. Robinson, Jad Farha, Amanda Bradke, Sherita H. Golden, Yanxun Xu, and Brian T. Garibaldi. Racial and Ethnic Discrepancy in Pulse Oximetry and Delayed Identification of Treatment Eligibility Among Patients With COVID-19. JAMA Internal Medicine, 182(7):730, July 2022. ISSN 2168-6106. doi: 10.1001/jamainternmed.2022.1906.

8. Kawaiola C Aoki, Maya Barrant, Mam Jarra Gai, Marina Handal, Vivian Xu, and Harvey N Mayrovitz. Impacts of Skin Color and Hypoxemia on Noninvasive Assessment of Peripheral Blood Oxygen Saturation: A Scoping Review. Cureus, September 2023. ISSN 2168-8184. doi: 10.7759/cureus.46078.

9. Michael W. Sjoding, Robert P. Dickson, Theodore J. Iwashyna, Steven E. Gay, and Thomas S. Valley. Racial Bias in Pulse Oximetry Measurement. New England Journal of Medicine, 383(25):2477–2478, December 2020. ISSN 0028-4793, 1533-4406. doi: 10.1056/NEJMc2029240.

10. Brandmaier. FDA Guidelines; Clinical Application of ARMS in Pulse Oximetry – Clinimark June 2021, 2018.

11. DIN EN ISO 80601-2-61: Medical electrical equipment - Part 2-61: Particular requirements for basic safety and essential performance of pulse oximeter equipment for medical use, 2017.

12. Chunhu Shi, Mark Goodall, Jo Dumville, James Hill, Gill Norman, Oliver Hamer, Andrew Clegg, Caroline Leigh Watkins, George Georgiou, Alexander Hodkinson, Catherine Elizabeth Lightbody, Paul Dark, and Nicky Cullum. The accuracy of pulse oximetry in measuring oxygen saturation by levels of skin pigmentation: a systematic review and meta-analysis. BMC medicine, 20(1):267, August 2022. ISSN 1741-7015. doi: 10.1186/s12916-022-02452-8.

13. Ralf E Harskamp, Luuk Bekker, Jelle C L Himmelreich, Lukas De Clercq, Evert P M Karregat, Mengalvio E Sleeswijk, and Wim A M Lucassen. Performance of popular pulse oximeters compared with simultaneous arterial oxygen saturation or clinical-grade pulse oximetry: a cross-sectional validation study in intensive care patients. BMJ Open Respiratory Research, 8(1):e000939, September 2021. ISSN 2052-4439. doi: 10.1136/bmjresp-2021-000939.

14. Nienke Bosschaart, Gerda J. Edelman, Maurice C. G. Aalders, Ton G. Van Leeuwen, and Dirk J. Faber. A literature review and novel theoretical approach on the optical properties of whole blood. Lasers in Medical Science, 29(2):453–479, March 2014. ISSN 0268-8921, 1435-604X. doi: 10.1007/s10103-013-1446-7.

15. Q. J. W. Milner and G. R. Mathews. An assessment of the accuracy of pulse oximeters. Anaesthesia, 67(4):396–401, April 2012. ISSN 0003-2409, 1365-2044. doi: 10.1111/j.1365-2044.2011.07021.x.

16. Mark S. Rea and Andrew Bierman. Light source spectra are the likely cause of systematic bias in pulse oximeter readings for individuals with darker skin pigmentation. British Journal of Anaesthesia, 131(4):e101–e103, October 2023. ISSN 00070912. doi: 10.1016/j.bja.2023.04.018.

17. Andrew Bierman, Kevin Benner, and Mark S. Rea. Melanin bias in pulse oximetry explained by light source spectral bandwidth. British Journal of Anaesthesia, 132(5):957–963, May 2024. ISSN 1471-6771. doi: 10.1016/j.bja.2024.01.037.

18. Raghda Al-Halawani, Meha Qassem, and Panicos A. Kyriacou. Monte Carlo simulation of the effect of melanin concentration on light–tissue interactions for transmittance pulse oximetry measurement. Journal of Biomedical Optics, 29(S3), August 2024. ISSN 1083-3668. doi: 10.1117/1.JBO.29.S3.S33305.

19. Andreia V. Moço, Sander Stuijk, and Gerard De Haan. New insights into the origin of remote PPG signals in visible light and infrared. Scientific Reports, 8(1):8501, May 2018. ISSN 2045-2322. doi: 10.1038/s41598-018-26068-2.

20. Lihong Wang, Steven L. Jacques, and Liqiong Zheng. MCML—Monte Carlo modeling of light transport in multi-layered tissues. Computer Methods and Programs in Biomedicine, 47(2):131–146, July 1995. ISSN 01692607. doi: 10.1016/0169-2607(95)01640-F. Number: 2.

21. Subhasri Chatterjee and Panayiotis Kyriacou. Monte Carlo Analysis of Optical Interactions in Reflectance and Transmittance Finger Photoplethysmography. Sensors, 19(4):789, February 2019. ISSN 1424-8220. doi: 10.3390/s19040789. Number: 4.

22. Olivier Tsiakaka, Benoit Gosselin, and Sylvain Feruglio. Source–Detector Spectral Pairing-Related Inaccuracies in Pulse Oximetry: Evaluation of the Wavelength Shift. Sensors, 20 (11):3302, June 2020. ISSN 1424-8220. doi: 10.3390/s20113302.

23. Thomas B. Fitzpatrick. The Validity and Practicality of Sun-Reactive Skin Types I Through VI. Archives of Dermatology, 124(6):869, June 1988. ISSN 0003-987X. doi: 10.1001/archderm.1988.01670060015008.

24. Maximilian Reiser, Timm Müller, Klaus Flock, Oliver Amft, and Andreas Breidenassel. Comparison of non-pulsating reflective PPG signals in skin phantom, wearable device prototype, and Monte Carlo simulations. In EMBC ‘23: Proceedings of the 45th Annual International Conference of the IEEE Engineering in Medicine & Biology Society, Sydney, Australia, December 2023. IEEE. doi: 10.1109/EMBC40787.2023.10340790.

25. M. Reiser, T. Mueller, A. Breidenassel, and O. Amft. Source-Detector Geometry Analysis of Reflective PPG by Measurements and Simulations. IEEE Open Journal of Engineering in Medicine and Biology, 6:400–406, 2025. ISSN 2644-1276. doi: 10.1109/OJEMB.2025.3546771.

26. Maximilian Reiser, Oliver Amft, and Andreas Breidenassel. Are VCSELs Better than LEDs for Wearable Reflective PPG? In Proceedings of the 2025 ACM International Symposium on Wearable Computers, pages 16–21, Espoo Finland, October 2025. ACM. ISBN 979-8-4007-1481-8. doi: 10.1145/3715071.3750423.

27. Maximilian Reiser, Andreas Breidenassel, and Oliver Amft. Impact of sensor configuration and melanin concentration on reflective pulse oximetry using Monte Carlo simulations. Scientific Reports, 15(1):39363, November 2025. ISSN 2045-2322. doi: 10.1038/s41598-025-26560-6.

28. Sophie Jenne. Flexible micro-spectrometers for agricultural and medical applications, 2024.

